# Myofibrillar Structural Variability Underlies Contractile Function in Stem Cell-Derived Cardiomyocytes

**DOI:** 10.1101/2020.10.13.336685

**Authors:** Kathryn Ufford, Sabrina Friedline, Zhaowen Tong, Vi T. Tang, Amani S. Dobbs, Yao-Chang Tsan, Stephanie L. Bielas, Allen P. Liu, Adam S. Helms

**Affiliations:** Division of Cardiovascular Medicine, University of Michigan; Department of Biophysics, University of Michigan; Department of Human Genetics, University of Michigan; Department of Neuroscience, University of Michigan; Department of Mechanical Engineering, University of Michigan; Department of Biomedical Engineering, University of Michigan, Ann Arbor, Michigan, USA; Cellular and Molecular Biology Program, University of Michigan, Ann Arbor, Michigan, USA

## Abstract

Disease modeling and pharmaceutical testing using cardiomyocytes derived from induced pluripotent stem cell (iPSC-CMs) requires accurate assessment of contractile function. Micropatterning iPSC-CMs on elastic substrates controls cell shape and alignment to enable contractile studies, but determinants of intrinsic variability in this system have been incompletely characterized. The objective of this study was to determine the impact of myofibrillar structure on contractile function in iPSC-CMs. Automated analysis of micropatterned iPSC-CMs labeled with a cell permeant F-actin dye revealed that myofibrillar abundance is widely variable among iPSC-CMs and strongly correlates with contractile function. This variability is not reduced by subcloning from single iPSCs and is independent of iPSC-CM purification method. Controlling for myofibrillar structure reduces false positive findings related to batch effect and improves sensitivity for pharmacologic testing and disease modeling. This analysis provides compelling evidence that myofibrillar structure should be assessed concurrently in studies investigating contractile function in iPSC-CMs.

## Introduction

Cardiomyocytes derived from induced pluripotent stem cell (iPSC-CMs) have high potential as a model system for investigating causal mechanisms in cardiomyopathies and for pharmaceutical testing. Multiple approaches have been reported to assess contractile function in iPSC-CMs, including motion assessment in 2D monolayers, 3D microtissue approaches, and the single cell micropatterning approach, as recently reviewed.(Blair and Pruitt, 2020) While 2D monolayer motion analysis is simple, disorganized myofibrillar organization limits accuracy and reproducibility. In contrast, Ribeiro et al. demonstrated that the single iPSC-CM micropatterning approach has advantages of single cell precision and control over cell shape, which was shown to exert a large influence on contractile function.(Ribeiro et al., 2015) Additionally, the single cell iPSC-CM method avoids interactions with non-myocytes, which are generally admixed into 3D iPSC-CM microtissues to maintain tissue integrity but may increase the potential for batch-to-batch heterogeneity.(Thavandiran et al., 2013)

Yet, despite the advantages of single cell precision and geometric constraint, substantial variability has remained in the single micropatterned iPSC-CM system.(Helms et al., 2020; Ribeiro et al., 2015) Since the method is time intensive, the number of replicates has been limited in early studies, and, consequently, the extent of variability in contractile parameters has not been characterized. Here, we develop a novel technique to perform unbiased quantification of myofibrillar structure in micropatterned iPSC-CMs using a live cell permeant dye that labels F-actin without requiring viral transduction or genetic engineering. Furthermore, using a large traction force microscopy (TFM) dataset of micropatterned iPSC-CMs, we construct a multivariable regression model that predicts contractile outputs from myofibrillar structural parameters. We present compelling evidence that variability in myofibrillar abundance is the key determinant of contractile function across individual iPSC-CMs. This variability is not due to genetic drift in heterogeneous stem cell populations, since it is not resolved by subcloning from single iPSCs; nor is this variability due to potential incomplete pluripotency in iPSCs, since the same observations extend to embryonic stem cell-derived cardiomyocytes (ESC-CMs). Through a power analysis, we show how this variability reduces sensitivity for differences between groups while simultaneously creating a tendency for false positive results by batch effect. Finally, we demonstrate how myofibrillar variability can mask substantial effects of pharmacologic contractile inhibition, and we apply the automated myofibrillar analysis method to an iPSC-CM cardiomyopathy model.

## Results

### Microcontact printing achieves highly consistent single iPSC-CM geometries but variable contractile function

To assess variability in micropatterned single iPSC-CMs, we selected the DF19-9-11 iPSC line (WiCell), a well-validated control cell line with similar calcium handling properties as other iPSC lines.(Hwang et al., 2015; Yu et al., 2009) We included iPSC-CMs purified by two alternative methods (selection by SIRPA membrane protein or metabolic selection with lactate-containing, glucose-deprived media) to assess influence on contractile assays.(Dubois et al., 2011; Tohyama et al., 2013) On unpatterned substrates, lactate-selected iPSC-CMs were larger with greater aspect ratios than SIRPA-sorted iPSC-CMs, but both conditions had broad variance (**Figure 1A-B**). In contrast, micropatterned iPSC-CMs with either method exhibited similar area and aspect ratio (**Figure 1A-B**). We analyzed contractile phenotypes in these micropatterned iPSC-CMs on 8.7 kPa hydrogels, simulating physiologic elasticity, at day 32 (**Figure 1C-D**). Although shape and eccentricity influence single cell contractile function,(Bray et al., 2008; Ribeiro et al., 2015) we discovered that marked variability was still evident in contractile function (**Figure 1E**).

**Figure 1.**
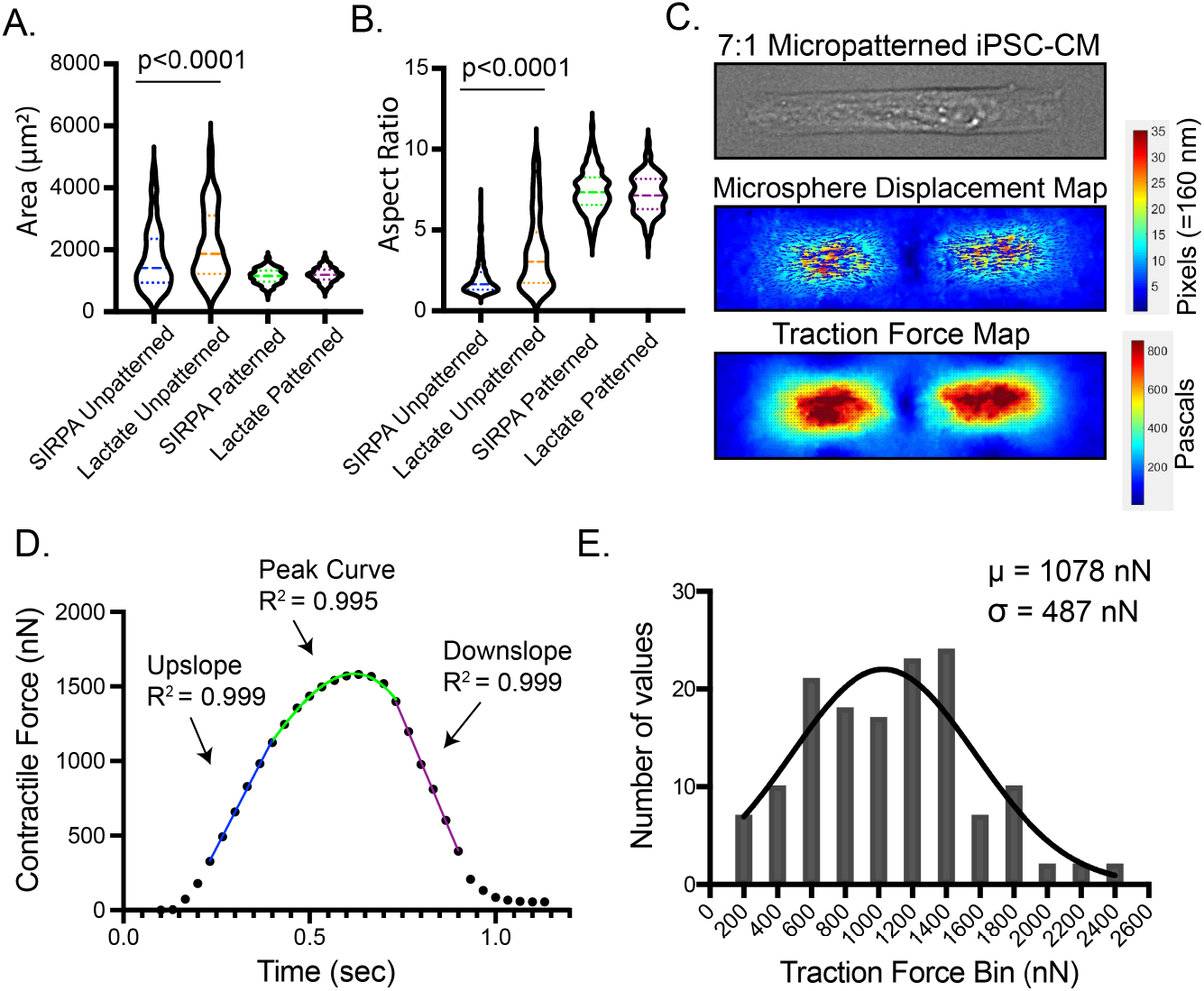
Microcontact printing achieves highly consistent single iPSC-CM geometries but variable contractile function. **A-B.** Area and aspect ratio of single iPSC-CMs both vary markedly with standard culture conditions but were reproducibly constrained with micropatterning (N=167 unpatterned SIRPA, N=121 unpatterned metabolic selection, N=53 micropatterned SIRPA, N=90 micropatterned lactate selection, comparisons by ANOVA). **C.** Representative displacement and traction force maps are shown at peak contraction. Displacements of microbeads were tracked and resolved to a traction force map to calculate the whole-cell traction force at 30 msec intervals. **D.** Representative analysis of a contractile curve demonstrates that partitioning into 3 components enables curve fitting to the calculate contraction velocity (upslope fit with linear regression, blue, invariably with R^2^>0.99), peak contraction (fit with polynomial, green, with typical R^2^>0.98), and relaxation velocity (fit with linear regression, red, invariably with R^2^>0.99). **E.** Histogram of maximum traction force across the population of individual micropatterned iPSC-CMs (N=143) demonstrates a Gaussian distribution with a standard deviation of 487 nN.

### Micropatterned iPSC-CMs exhibit reproducible myofibrillar alignment but variable myofibrillar abundance

The consistency and variability of myofibrillar development across large populations of micropatterned iPSC-CMs has not been reported but may underlie the variability in contractile function. To characterize myofibrillar structure systematically, we performed sequential live imaging of F-actin using a cell permeant dye (SiR-actin) following traction force microscopy to obtain near-simultaneous images (**Figure 2**). We then performed an unbiased quantification of myofibrillar structure using an automated pipeline in MATLAB^®^ that outputs total cell area, aspect ratio, myofibrillar bundle area, density, alignment, and heterogeneity (available at https://github.com/Cardiomyocyte-Imaging-Analysis/MyofiberQuant). Using this approach, we identified highly reproducible alignment of myofibrils but marked variability in myofibrillar structure across iPSC-CMs (**Figure 2A-D**).

**Figure 2.**
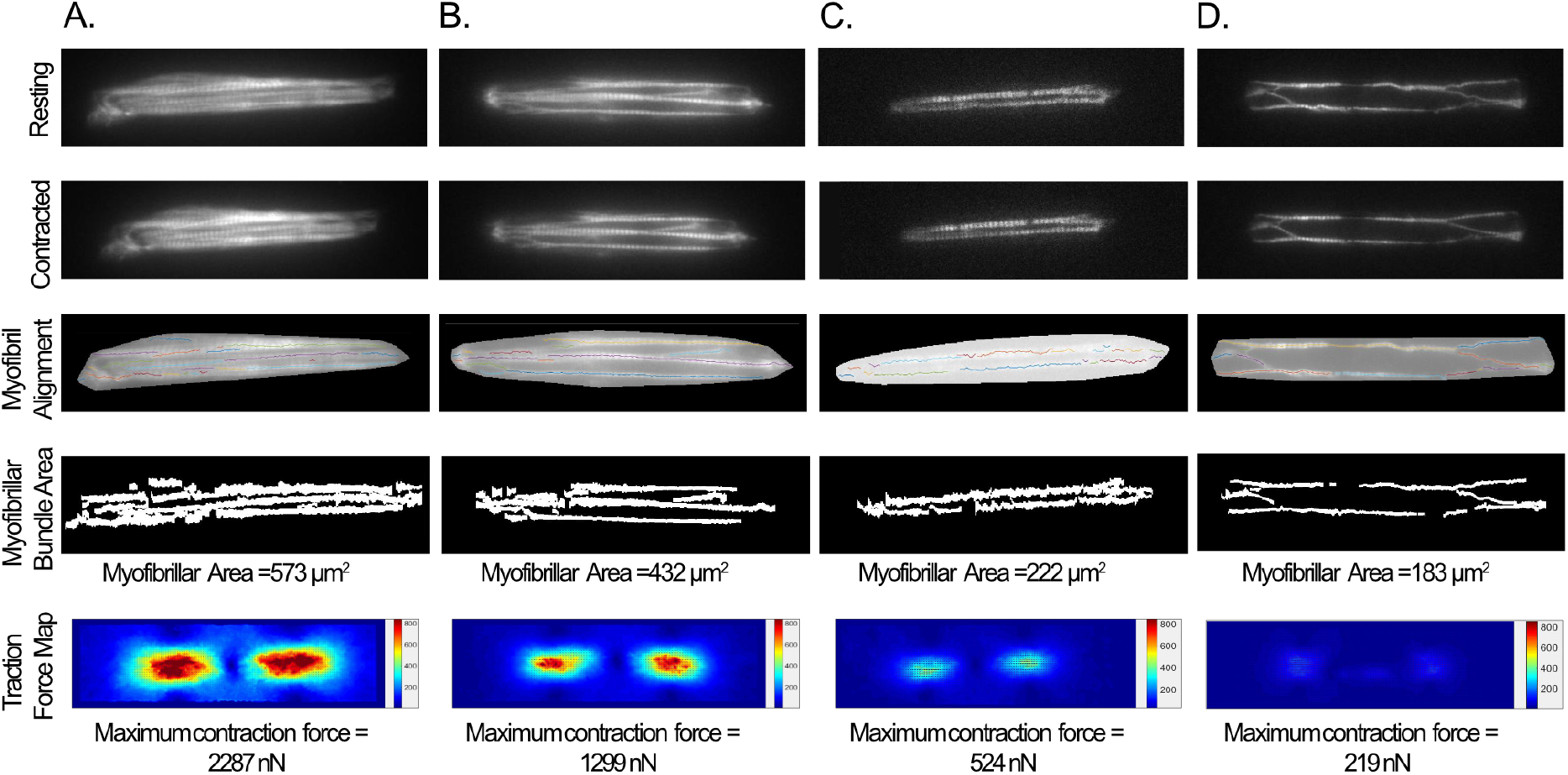
Single micropatterned iPSC-CMs demonstrate consistent myofibrillar alignment but variable myofibrillar structure and abundance. **A.** Representative images of a micropatterned iPSC-CM with dense myofibrillar bundles (fluorescence imaging of F-actin) obtained at time of traction force microscopy. Time series of images of F-actin were collected throughout the contraction cycle to avoid blur (top row at rest; 2^nd^ row at peak contraction). An automated image processing algorithm was used to extract a masked cell area and identify myofibrillar bundles. The algorithm calculates myofibrillar alignment (3^rd^ row) and total myofibrillar bundle area (4^th^ row), as well as heterogeneity in both short- and long-axis directions. A corresponding traction force map is shown in the 5^th^ row. **B-D**. Representative images of micropatterned iPSC-CMs across a range of myofibrillar densities with corresponding traction force maps. Myofibrillar alignment was robust along the long axis for all iPSC-CMs, but contractile force was diminished in iPSC-CMs with low myofibrillar density (C-D).

### Myofibrillar structural parameters predict contractile function in micropatterned iPSC-CMs

We hypothesized that the observed variability in contractile phenotypes is due to differential myofibrillar structure in individual iPSC-CMs. To test this hypothesis, we generated a dataset of traction force microscopy and myofibrillar structural measurements for 12 batches of control iPSC-CMs. With linear regression, myofibrillar area significantly correlated with contractile force (**Figure 3A**, R^2^ = 0.32, p<0.0001). Among other structural parameters, total cell area and heterogeneity index showed modest correlations with contractile force (R^2^ = 0.10, R^2^ = 0.05; **Figure S1**). Construction of a multivariable model of all structural parameters improved model prediction of contractile output compared to simple regression with myofibrillar area alone (**Figure 3B and S2**, R^2^ = 0.43, p<0.0001 vs simple regression). In the multivariate model, only myofibrillar area was independently and significantly associated with contractile force (p<0.0001).

**Figure 3.**
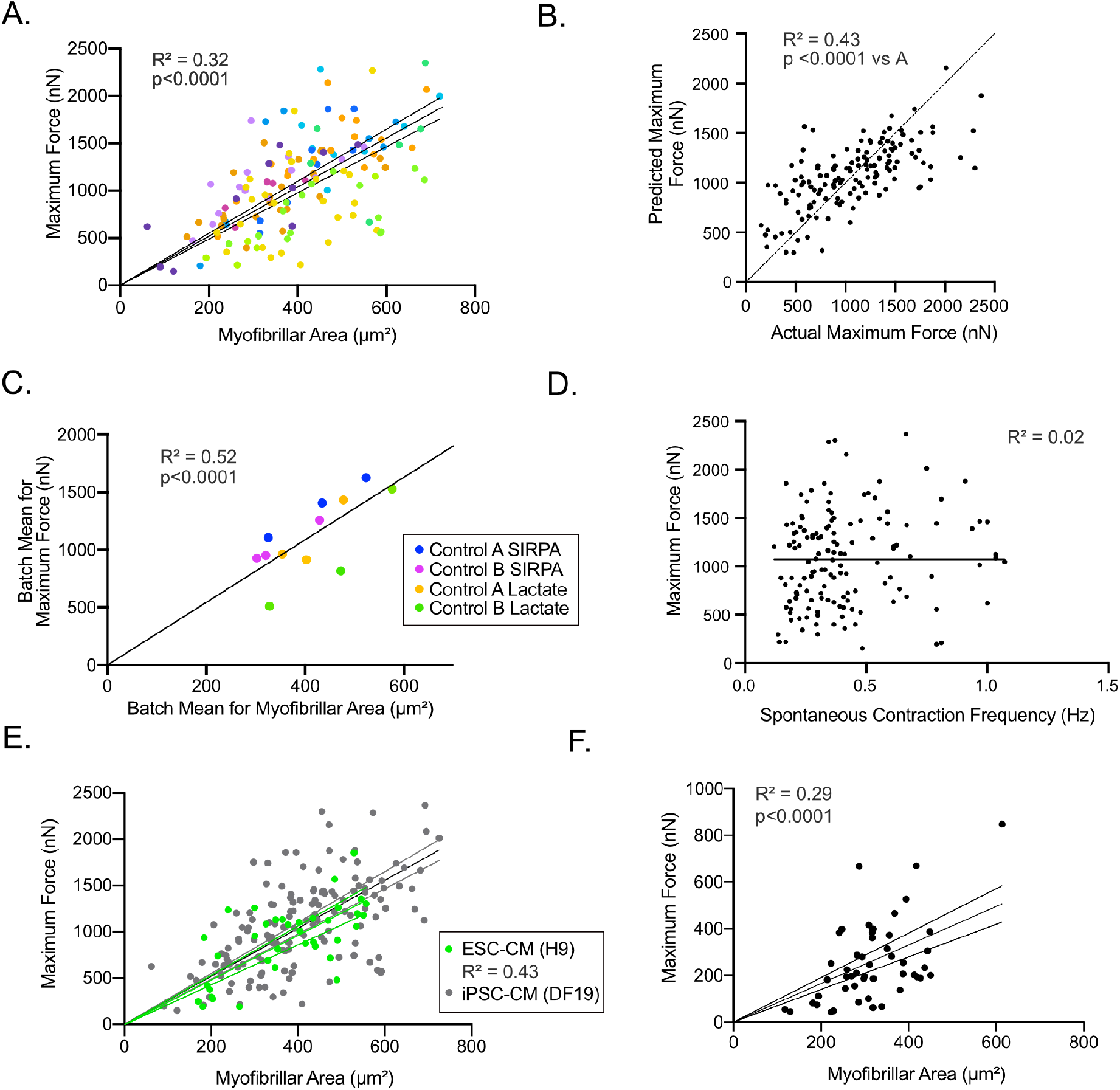
Myofibrillar structure correlates with maximal contraction force. **A.** Linear regression demonstrates a significantly positive correlation between myofibrillar area and maximum contraction force (N=143 from 12 batches of iPSC-CMs, p<0.0001 vs. slope of 0, trend line with 95% confidence interval regression lines shown). Each iPSC-CM batch is shown as a different color (shades of corresponding colors match batch colors in 2D). **B.** Actual versus predicted maximum traction force from multiple regression incorporating myofibrillar area, cell area, aspect ratio, and heterogeneity indices. **C.** Mean myofibrillar area per batch correlated with mean contractile force across control iPSC-CM batches for both control lines purified by either SIRPA sorting or metabolic selection. **D.** Spontaneous contraction frequency does not correlate with contractile force. **E.** Linear regression of myofibrillar area versus maximum contractile force for ESC-CMs (green points and regression line +/- 95% confidence intervals, N=37, p<0.0001 vs. slope of 0) demonstrated a similar relationship as for iPSC-CMs. ESC-CM points are overlayed on iPSC-CM points for comparison (iPSC-CMs shown as gray points and regression line +/- 95% confidence intervals, N=143, same as in Figure 3A). **F.** iPSC-CMs plated on substrates with higher elastic modulus (25 kPa) also demonstrated a significant linear correlation between myofibrillar abundance and contractile force (N=49 from 3 batches, p<0.0001 vs. slope of 0). See also Figures S1-S2.

### Myofibrillar structural parameters associate with batch variability in contractile function in micropatterned iPSC-CMs

We next examined whether variability in myofibrillar abundance could be associated with batch-to-batch variability in contractile function. Linear regression of mean values for each batch demonstrated that myofibrillar area strongly correlated with mean contractile force for each batch (R^2^ = 0.52, p<0.0001, **Figure 3C**). Control iPSC-CM batches consisted of 6 each from the DF19-9-11 control line and a single-cell derived subclone from this line (obtained as a non-targeted control line from a prior gene-editing experiment).(Helms et al., 2020) We observed similar variability in both myofibrillar and contractile phenotypes in both the originating line (control A) and subclone (control B) iPSC-CMs across cells and batches (**Figure 3A, 3C**). This result indicates that variability in myofibrillar and contractile phenotypes arise primarily from the differentiation environment and are not primarily due to genetic drift of individual iPSCs prior to differentiation. The dataset also included an equal number of batches of SIRPA or lactate-purified iPSC-CMs (6 each, equally divided among control A and control B) to assess any potential impact of purification method on contractile force variability. Contractile force was similar between purification methods (**Figure S1**), and myofibrillar abundance and contractile force were similarly correlated independent of purification method (**Figure 3C and S1**).

### Calcium transient quantification but not contraction frequency further improves multivariable prediction of contractile function

Since contraction frequency correlates with calcium transient amplitudes and contractile force in the heart, we assessed whether spontaneous contraction frequency of iPSC-CMs contributes to variability of contractile function. No significant correlation was observed (R^2^ = 0.02, **Figure 3D**). This finding is likely due to the low contraction frequency of iPSC-CMs in this study (75% <0.5 Hz, 96% <1.0 Hz), since the frequency-force response primarily exists at >1.0 Hz.(Endoh, 2004) Inclusion of frequency in the multivariable model did not improve fit (p=0.53).

To test whether residual variability in contractile function could be due to variability in calcium transients independent of frequency, we quantified transient amplitudes in a subset of iPSC-CMs. In simple linear regression, calcium transient amplitude did not correlate with force (**Figure S3**). However, inclusion of transient amplitude in a multivariable analysis resulted in an improved model fit compared to regression with myofibrillar area alone (**Figure S3**, R^2^ = 0.60 vs. R^2^ = 0.33, p=0.002). Thus, while myofibrillar abundance most strongly correlates with contractile function, variability in calcium handling additionally contributes to contractile phenotypes across iPSC-CMs.

### The myofibrillar area and contractile function relationship extends to embryonic stem cell-derived cardiomyocytes and is independent of culture substrate stiffness

Although the DF19-9-11 iPSC line has been widely used, we further tested whether potential incomplete pluripotency in this line could explain the variability in myofibrillar development by performing the same quantifications using an embryonic stem cell (ESC) line. Differentiated cardiomyocytes from the H9 ESC line (ESC-CMs) also exhibited marked variability, and structure-function relationships for individual H9 ESC-CMs distributed similarly as iPSC-CMs with a similar regression of myofibrillar area vs. force (**Figure 3E**). We also tested whether the cell culture substrate stiffness might impact the relationship between myofibrillar structure and contractile function. On 25 kPa stiff substrates, we also observed a correlation between myofibrillar abundance and contractile force in iPSC-CMs (**Figure 3F**).

### Concurrent quantification of myofibrillar structure reduces false positive results and improves sensitivity of iPSC-CM contractile studies

Although statistical tests generally used for iPSC-CM contractility studies assume an unbiased distribution of phenotypes across batches, batch-related bias is a generally acknowledged limitation and could result in false positive findings. Since a practical number of batches (often 3) is typically chosen, we assessed the potential for false positive results related to batch bias by comparing all possible combinations of comparison groups of 3 from the 6 SIRPA-purified control batches in our dataset. Out of 10 possible combinations, 3 comparisons of contractile force were statistically significant, consistent with a 30% false positive rate (**Figure 4A**). Normalization of contractile force by myofibrillar area prevented these false positives (**Figure 4B**).

**Figure 4.**
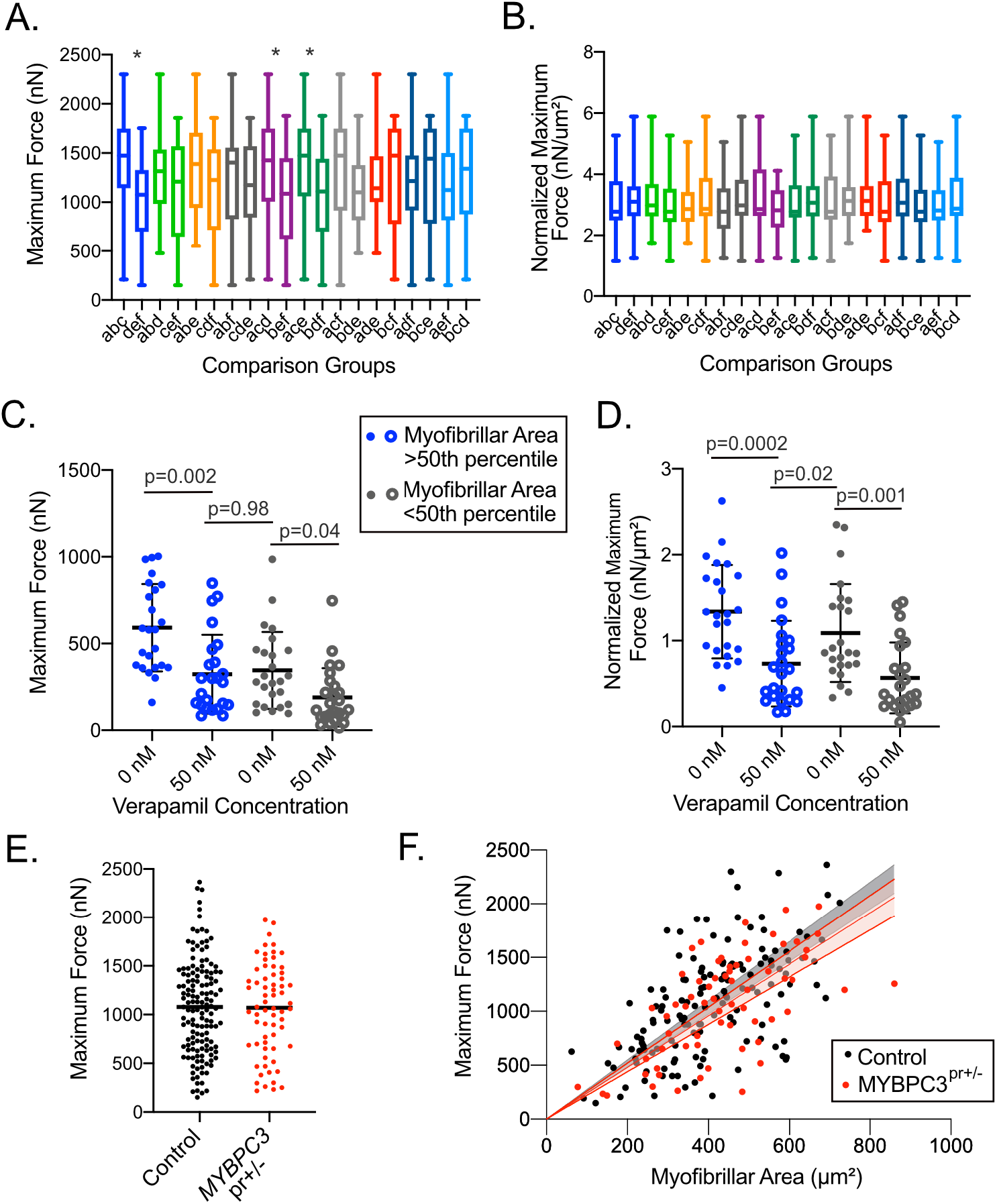
Concurrent quantification of myofibrillar structure reduces false positive results and improves sensitivity of iPSC-CM contractile studies. **A.** The 6 control SIRPA-selected batches from Figure 3 were subdivided into all possible combinations of 3 batches vs. 3 batches to assess the influence of batch effect on statistical comparisons of contractile force. Each unique 2-group combination is shown in a different color and were compared using the independent samples t-test (box plot shows 75% interquartile with whiskers showing range; * indicates p<0.05). **B.** Normalization of contractile force by myofibrillar area results in a lack of statistically significant differences among the same groups as in A. **C.** The effect of myofibrillar area on quantification of contractile force response to verapamil was assessed with iPSC-CMs stratified by myofibrillar abundance. Within the stratified groups, verapamil reduction of contractile force was statistically significant but comparison across groups with different myofibrillar abundance precluded detection of reduced contractile force. **D.** Normalization of contractile force by myofibrillar area per iPSC-CM enabled discrimination of iPSC-CMs treated with verapamil irrespective of myofibrillar abundance (for C-D, total N=49, p<0.05 by ANOVA with Sidak’s multiple testing correction considered significant). **E.** Contractile measurements in both control and *MYBPC3*^pr+/-^ iPSC-CMs exhibit marked variability (coefficient of variation 0.45 and 0.44, respectively). **F.** Linear regression demonstrates a similar relationship between myofibrillar area and contractile force for MYBPC3 ^pr+/-^ iPSC-CMs (shaded 95% confidence intervals of regression lines). Multiple regression including genotype did not improve model fit (p=0.36, extra sum-of-squares F test).

Next, we assessed the sensitivity of single cell iPSC-CM contractile studies by performing a power analysis using the standard deviation of traction forces (σ=487 nN) from the control cell population (**Figure 1E**). To identify a 10% difference in maximum contractile force with β (type II error) = 0.9 and α (type I error) = 0.05, a sample size of 974 cells is required for a 2-group comparison (487 cells per group) – far surpassing the number of replicates in early studies using these techniques.

We assessed the impact of myofibrillar variability on pharmacologic analyses by quantifying paired iPSC-CMs before and after 50 nM verapamil, which reduced mean contractile force by 42% (419 ± 251 nN vs. 242 ± 195 nN, p<0.0001). IPSC-CMs were stratified into two groups based on myofibrillar abundance greater or less than the 50^th^ percentile. We found that the group of iPSC-CMs with high myofibrillar abundance exerted similar contractile force post-verapamil as iPSC-CMs with low myofibrillar abundance pre-verapamil (**Figure 4C**). This result shows that even a marked difference in contractility can be masked by differences in myofibrillar structure. Normalization of contractile force by myofibrillar area revealed a significant reduction irrespective of myofibrillar abundance (**Figure 4D**). Together, these data show that batch effect increases the likelihood of false positive results from iPSC-CM contractile studies while broad intrinsic variability reduces the sensitivity for actual differences. Controlling for myofibrillar structure both reduces false positive findings and improves sensitivity.

Finally, we utilized this analysis method for a cardiomyopathy iPSC-CM model with heterozygous knock-out of the sarcomere gene *MYBPC3* via promoter deletion (*MYBPC3*^pr+/-^) that we previously reported, since effects of heterozygous *MYBPC3* loss of function mutations on maximal contraction force have been inconsistent across iPSC-CM studies.(Helms et al., 2020) In the prior study, we used a simple binary mask threshold to select cells above a threshold of 40% myofibrillar density for analysis, potentially creating selection bias. Here, we employed the automated analysis pipeline to an expanded set of all imaged cells. Similar to controls, *MYBPC3*^pr+/-^ iPSC-CMs exhibited broad variability in contractile force (**Figure 4E**). Multiple regression stratified by genotype confirmed a lack of difference in contractile force in *MYBPC3*^pr+/-^ iPSC-CMs (**Figure 4F**).

## Discussion

Micropatterning enables precise geometric control for single cell biomechanical studies. In this study, we used a technique that creates μm-level precision of adhesive protein on polyacrylamide gels formulated for a physiologic workload. Similar to results with neonatal rat ventricular myocytes and iPSC-CMs using similar techniques(Bray et al., 2008; Ribeiro et al., 2015), we found that iPSC-CMs adhere to micropatterns with ~7:1 aspect ratios, inducing alignment of myofibrils. Yet, despite robust consistency of shape and alignment, marked variability in contractile function persisted. Our analysis reveals that this variability is largely due to underlying variability of myofibrillar abundance across individual iPSC-CMs. This variability creates a major obstacle to reproducible functional measurements and requires unbiased methods for rigorous assessment.

The cause of variable myofibrillar development in micropatterned iPSC-CMs could be differential exogenous signaling cues in the preceding monolayer environment, including influences from mixed cell types. To reduce these factors, we purified iPSC-CMs using two different methods, both yielding similar results, and replated iPSC-CMs as purified monolayers prior to seeding iPSC-CMs to micropatterns. The persistence of variability may point to a stalling in maturation that is heterogeneous across populations of cells. We found that a longer culture time (>32 days) did not result in a significant improvement in contractile force (not shown). Wheelwright et al. observed that most improvement in contractile function occurred from day 14 to day 30, with only a modest improvement with extension to 90 days.(Wheelwright et al., 2018) Our findings are consistent with single cell RNA-seq analyses that demonstrated stalled and variable development in iPSC-CMs. (Friedman et al., 2018) Given this marked heterogeneity, assessment of myofibrillar structure through an unbiased approach enables verification of similarity among comparison groups. The automated technique developed here could be used to reduce the sample size in iPSC-CM studies, improve sensitivity, and reduce the potential for false positive results related to batch-to-batch variability.

Myofibrillar structural parameters explained a large degree, but not all, of variability in contractile function. One likely explanation for residual variability is that a 2-dimensional approach for quantifying myofibrillar abundance underestimates myofibrils in regions of high myofibrillar density due to a limited resolution of superimposed structures. Z-stack imaging with standardized absolute intensity quantification may increase accuracy but would not have enabled live cell contractile imaging. Additionally, we performed a linear regression based on the assumption that myofilament addition in parallel would be linearly related to contractile output. Actual intracellular relationships among myofilament organization and contractile output are likely more complex. Finally, contractile function may also vary due to cell-cell differences in transcriptional programs independent from myofibrillar abundance, such as sarcomere gene transcript variant / isoform expression (e.g. TNNI3) and metabolic capacity (Birket et al., 2015; Correia et al., 2017; Wheelwright et al., 2020).

In conclusion, our results are compelling evidence that single micropatterned iPSC-CMs exhibit marked myofibrillar structural variability that underlies variability in contractile function. This variability is not resolved by standard purification methods or by subcloning to reduce genetic heterogeneity. Simultaneous imaging with an F-actin cell-permeant dye and an automated imaging analysis pipeline allow concurrent assessment of myofibrillar structure to verify similarity of comparison groups, improve sensitivity, and reduce false positive results. Moreover, future efforts should be directed toward reducing developmental variability in differentiated iPSC-CMs.

## Experimental Procedures

Stem Cell and Cardiomyocyte Culture: Control iPSCs (DF19-9-11) and ESCs (H9 WA09) were obtained from WiCell. An isogenic, single cell derived subclone of the DF19-9-11 line and the isogenic MYBPC3^pr+/-^ line were derived during a previously reported gene-editing experiment.(Helms et al., 2020) IPSCs were cultured in mTeSR1 and differentiated using Wnt modulation.(Lian et al., 2012) IPSCs were verified free of mycoplasma using MycoAlertTM (Lonza). iPSC-CM were purified by either SIRPA-selection using magnetic beads at day 15 (Herron et al., 2016) or metabolic selection using glucose-deprived, lactate-containing media on days 12-16.(Burridge et al., 2015; Tohyama et al., 2013) Purified iPSC-CMs were first replated as monolayers until day 25, then replated to micropatterned substrates at 6000k/cm^2^ density until assay at day 32. Because RPMI contains a subphysiologic level of calcium (0.4 mM), RPMI was blended 50:50 with DMEM (1.10 mM calcium) on day 1 following replating, and then 25:75 with DMEM (1.45 mM calcium) on day 3 following single cell replating (1X B27 supplement).

Microcontact Printing and Live-Cell Imaging: Single cell micropatterning was performed as described.(Helms et al., 2020) A chrome mask with grids of 7:1 aspect ratio, 1750 μm^2^ area rectangles was used to generate a silicon wafer with micropatterned spin-coated SU-8. PDMS stamps were molded with Sylgard^®^ 184 (Corning). Polyacrylamide hydrogels were fabricated onto glass coverslips at either 8.7 kPa or 25 kPa by altering the proportion of bis-acrylamide, and microprinted with 1:25 diluted 0.1% human serum-derived fibronectin (Sigma).(Grevesse et al., 2013, 2014; Helms et al., 2020) Hydrogels contained 0.2 μm diameter FluoSpheresTM (Thermo Fisher). All iPSC-CMs were labeled for F-actin with SiR-Actin (Cytoskeleton, Inc.), prior to imaging (0.5 μM, 1 hour). Medias were equilibrated (37° C and 5% CO_2_) prior to media changes. Microscopy was performed at 37° C and 5% CO_2_ to acquire videos of spontaneously contracting iPSC-CMs, fluorescent microbeads, and myofibrils. Only iPSC-CMs exhibiting spontaneous contractions with both ends symmetrically contracting toward the center were included.

Image Processing: Traction force analysis was performed as described (see Supplemental Information).(Han et al., 2015; Helms et al., 2020) Images of myofibrils were processed using custom scripts in MATLAB^®^ that extract intensity-based signals for individual myofibrillar bundles (see Supplementary Information, available on GitHub).

Intracellular Calcium Quantification: iPSC-CMs were loaded with 1 μM fura-2 AM (Thermo Fisher) for 10 min, rinsed with 250 μM probenecid in HBSS for 5 min, changed to pre-equilibrated media, and then incubated for 30 min prior to imaging through a Nikon Super Fluor 40x objective. The emitted light intensity following alternating excitation at 340 nm and 380 nm was reported as fura-2 ratios.

Statistical Analysis: Continuous variables were compared by ANOVA with multiple testing correction. Linear regressions were performed with the least squares method. Goodness of fit was assessed by R^2^, and models were compared with the extra-sum-of-squares F-test. All analyses were performed in GraphPad Prism.

## Acknowledgements

This work was funded by the NIH (K08HL130455 to ASH and R21GM134167 to APL) and NSF (CMMI-1561794 to APL). We thank Kevin Kit Parker and Thomas Grevesse for initial training in polyacrylamide gel micropatterning, Sangyoon Han for assistance in implementation of traction force analysis software, and Lap Man Lee for fabrication of the micropatterning wafer.

## Author Contributions

Conceptualization - ASH, APL; Methodology - ASH, APL; Investigation - KU, SF, ZT, VTT, AD, YCT; Writing - ASH, KU; Writing, Review/Editing - ASH, APL, SLB; Funding – ASH, APL; Supervision – ASH, APL, SLB.

