## Supplemental Information for "Myofibrillar Structural Variability Underlies Contractile Function in Stem Cell-Derived Cardiomyocytes"

Ufford, et al.

### Supplemental Figures

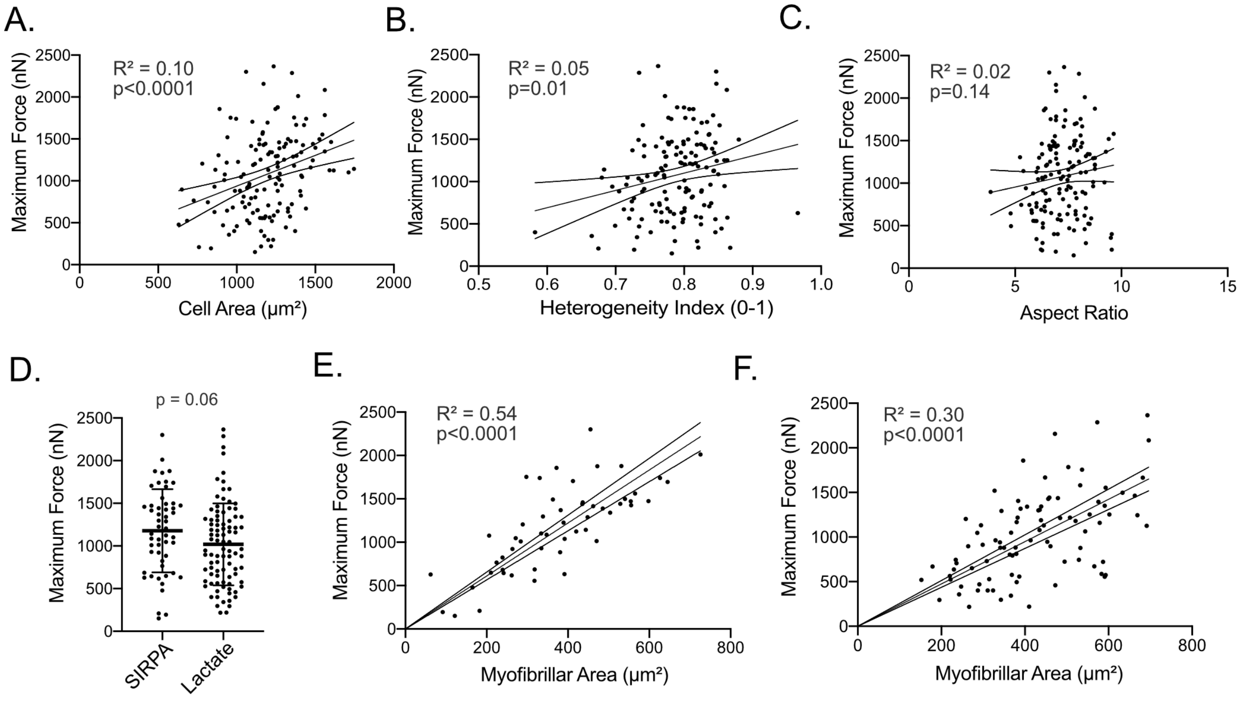


**Figure S1. Univariate analyses of structural parameters and purification method with iPSC-CM contractile force.** Related to Figure 3. **A-C.** Among micropatterned iPSC-CMs, simple linear regression analysis showed a modest correlation between cell size and contractile force (A) and between heterogeneity index and contractile force (B) but no significant correlation for aspect ratio. **D.** Univariate analysis showed no statistically significant association between iPSC-CM purification method and contractile force. **E-F.** When analyzed separately with simple linear regression (as compared to Figure 3A), subsets of iPSC-CMs purified by SIRPA selection (E, N=53) and by lactate selection (F, N=90) independently demonstrated a significant correlation between myofibrillar area and contractile force. Goodness of fit for linear regressions was assessed by $R^2$, and the linear slope of each regression was compared to a slope of 0 by the extra sum-of-squares F-test ($p<0.05$ considered significant).

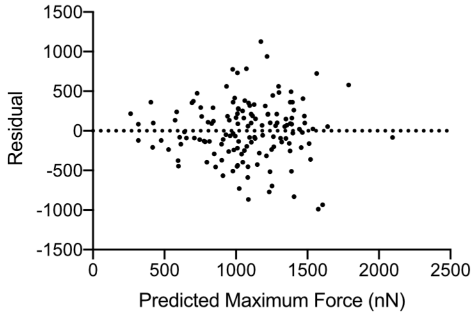


**Figure S2. Multivariate analysis residuals.** Related to Figure 3. The residual contractile force not predicted by the multivariate model is shown.

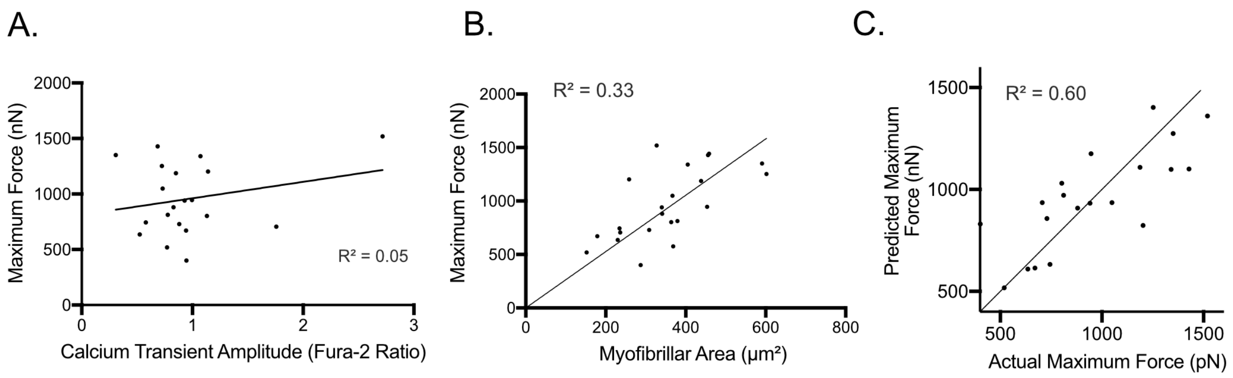


**Figure S3. Calcium handling contributes to contractile force prediction.** Related to Figure 3. **A.** Calcium transient amplitudes did not correlate with contractile force with simple linear regression (N=20). **B.** Simple linear regression showed a correlation between myofibrillar area and maximum contractile force for this subset of cells (N=20), similar to the larger population of iPSC-CMs shown in Figure 3A. **C.** Addition of calcium transient amplitudes to myofibrillar area in a multivariable model improved model fit (N=20, $R^2$ = 0.60 vs. $R^2$ = 0.33, p=0.002).

**Supplemental Experimental Procedures**

Traction Force Microscopy Analysis

Traction forces for each cell were quantified from time series imaging of iPSC-CMs using the method of Han, et al.(Han et al., 2015) Program settings for displacement field calculations included high-resolution subsampling of beads, and no outward deformation was expected. The displacement field was corrected for outliers at the strictest setting (“1”). The force field was calculated for a Young’s modulus of either 8.7 kPa or 25 kPa as relevant and gel thickness of 52 µm. Forces were reconstructed with the FastBEM method. The L-curve for regularization was not necessary given that standard imaging parameters were used throughout experiments. Mesh points were set to 4096 and the “backslash” coefficient solving method was used. Traction forces were calculated over the whole field of view and time-force data were exported to Microsoft Excel® for calculation of contractile kinetic parameters.

Myofibrillar Quantification Algorithm

Images of myofibrils were processed using custom scripts in MATLAB® that extract intensity-based signals for individual myofibrillar bundles by identification and merger of F-actin signal peaks connected along the long axis of the image. A region of interest which included the cell was first determined by separating individual pixels into two clusters based on intensity using k-means grouping. The pixels in the cluster with the higher intensity were identified as the foreground. These pixels were dilated, small holes were filled in, and the edges were smoothed to identify the cell boundary. After the region of interest was determined, the major axis of the cell was identified. The region of the image containing the cell had the original intensity and areas outside the cell were set to zero.

For the entire length of the cell, the image intensity along a vertical cross-section of the cell was plotted. Peaks, which could potentially correspond to myofibrils, were identified from this plot based on criteria including the height and width of the peak. To lessen the impact of signal from adjacent myofibrils, peaks were identified starting from the outer edge of the cell (where there was less signal amplification from neighboring myofibrils). After identification of each peak, the intensity plot of the peak was fit with a Gaussian curve. The width of the myofibril at that location was set equal to the standard deviation of the Gaussian fit.

After peaks potentially corresponding to myofibrils had been identified across the length of the cell, peaks related to the same myofibril were identified. This was done by starting with an identified peak and determining if there was another pixel in its neighborhood. The neighborhood was determined as the area within a radius of five pixels from the initial peak. If there was a peak within that neighborhood, the two peaks were connected and identified as being part of the same myofibril. If there was not a peak within that neighborhood, the peak was ignored in further steps. From there, the ending point of the myofibril was identified as the peak which was found within the neighborhood and the process was repeated across the length of the cell with myofibrils growing in length. Myofibrils which were within four pixels of one another and had similar trajectories were merged to become one longer myofibril. The orientation of each myofibril relative to the long axis of the cell was then determined. The total area containing myofibrils and the myofibrillar density were then calculated. Heterogeneity of myofibrillar distribution was calculated as an index (0-1) for which 1 indicates completely homogeneous distribution of myofibrils across the long-axis of the cell. Outputs of the algorithm include: cell area, myofibrillar area, myofibrillar density, heterogeneity index, and myofibrillar alignment.
